# Born Connected: The Early Emergence of Adult-Like Multi-Scale Brain Networks in Infancy

**DOI:** 10.1101/2024.11.27.625681

**Authors:** Prerana Bajracharya, Zening Fu, Shiva Mirzaeian, Vince Calhoun, Sarah Shultz, Armin Iraji

**Affiliations:** Tri-Institutional Center for Translational Research in Neuroimaging and Data Science (Georgia State University, Georgia Institute of Technology, Emory University), Atlanta, GA 30303, USA; Department of Computer Science, Georgia State University, Atlanta, GA 30303, USA; Division of Autism & Related Disabilities, Department of Pediatrics, Emory University School of Medicine, Atlanta, GA 30322, USA; Marcus Autism Center, Children’s Healthcare of Atlanta, Atlanta, GA, USA

**Author notes:** Correspondence to: Name: Prerana Bajracharya, 55 Park Place NE, 18th Floor – TReNDS Center Atlanta, GA – 30303, Name: Armin Iraji, 55 Park Place NE, 18th Floor – TReNDS Center Atlanta, GA – 30303.

**Keywords:** infant, intrinsic connectivity networks, functional network connectivity, multi-scale, burstICA, NeuroMark framework

## Abstract

The human brain undergoes remarkable development during the first six postnatal months, a period of dramatic structural and functional change critical for understanding neurodevelopmental trajectories. Previous infant resting-state functional MRI (rsfMRI) studies identified intrinsic connectivity networks (ICNs) but reported widely varying network numbers and organization, limiting cross-study comparisons. A recent analysis of over 100,000 subjects spanning from adolescents to adults generated a 105-network multi-scale template that greatly enhanced replicability, but the presence of these canonical ICNs in infants has not been investigated.

We analysed resting-state scans from infants aged 0–6 months using two complementary approaches: burst independent component analysis (burstICA), a fully data-driven, model-order-agnostic method, and the reference-informed NeuroMark framework. burstICA successfully recovered the full set of canonical ICNs directly from infant data, and simulation-based validation confirmed that the recovered networks were biologically specific rather than methodological biases. The NeuroMark framework enhanced spatial correspondence with the reference template and maintained distinct, reproducible network profiles across scans, demonstrating high reliability for cross-study and cross-age comparisons.

Networks critical for higher-order cognition, such as the default mode and salience networks, displayed adult-like topography comparable to that of primary sensory networks even within the first 6 months after birth. Subcortical networks also exhibited precise spatial organization, underscoring early maturation of deep brain structures. Functional network connectivity analyses revealed close similarity to adult profiles and clear antagonistic patterns between sensory systems and higher-order networks, indicating that foundational aspects of mature brain organization are already emerging in early infancy.

Together, these findings establish a methodological foundation for early network mapping in infancy and lay critical groundwork for longitudinal and translational studies of brain development.

## Introduction

Brain function is widely characterized through the principles of functional segregation and integration^1^, wherein the brain comprises distinct functional sources. In resting-state fMRI, these sources are estimated as intrinsic connectivity networks (ICNs), spatially distinct patterns of synchronized activity that support emerging cognitive and behavioral capacities^2^. Although the brain at birth is relatively immature, infancy is marked by rapid maturation in both anatomy and functional connectivity^3–5^, during which these ICNs are expected to evolve toward their mature configurations. Despite their central role in early brain development, the study of ICNs in infancy remains comparatively nascent relative to the adult literature.

Although still an emerging field, research on ICNs in infancy has begun to reveal consistent patterns of early functional brain organization. Initial studies, such as those by Liu et al.^6^ identified distinct sensorimotor networks in full-term neonates, including unilateral networks associated with hand movements and a midline network linked to lower limb and supplementary motor areas^6^. Gao et al.^7^ extended this work by mapping the developmental trajectories of 9 resting-state networks in infants from birth to one year, finding that primary networks, such as sensorimotor and auditory, are largely established at birth, while visual networks mature shortly thereafter^7^. In contrast, higher-order networks, including the default mode, dorsal attention, salience, and frontoparietal networks, exhibit more protracted development across the first year, with the salience and frontoparietal networks showing the slowest maturation^7^. More recent studies using the developing Human Connectome Project (dHCP) dataset, such as that by Eyre et al.^8^, identified 11 ICNs in term-born infants, distinguishing between early-developing primary networks (medial motor, lateral motor, somatosensory, auditory, visual) and later-developing association networks (motor association, temporoparietal, posterior parietal, frontoparietal, prefrontal, visual association)^8^. Similarly, Molloy and Saygin reported the presence of several adult-like ICNs, including sensorimotor, visual, default mode, ventral attention, and dorsal attention networks, in neonates, while noting the absence of limbic and control networks, indicating ongoing specialization^9^. Collectively, these studies highlight both the early emergence and continued maturation of functional networks during infancy, with primary sensory and motor networks emerging first, and higher-order association networks developing more gradually.

Despite these advances, significant gaps and challenges remain in the study of infant ICNs. One critical issue is the variability in the number of networks identified across studies, with some reporting as few as five networks^9^, while others detect up to eleven distinct networks^8^. This inconsistency poses significant challenges for cross-study comparisons and hinders efforts to establish reliable longitudinal trajectories of brain network development in early life. Additionally, many studies focus selectively on well-known networks, such as the language^10^ or default mode networks^11,12^, often overlooking other essential ICNs, resulting in a fragmented understanding of network evolution. This bias limits insights into the comprehensive development of brain connectivity in infancy, potentially missing the co-development of complex networks alongside primary ones. Furthermore, only few studies directly compare infant ICNs to those of mature adult samples^9^, leaving open questions about whether observed infant networks are early forms of adult-like ICNs or are transient configurations unique to infancy.

Another major gap in current infant neuroimaging research is the lack of multi-scale analyses. Most existing studies have relied on a single set of regions of interest or fixed-model independent component analysis (ICA)^6–8,11,13,14^, thereby constraining investigations to a single spatial scale. However, functional sources like ICNs are inherently organized across multiple spatial scales, where spatial scale refers to the level of granularity at which functional organization is resolved^15–17^. Large-scale networks, typically captured at low ICA model orders^18^, reflect broad and integrated systems, for example, an undivided default mode network, whole-brain sensorimotor network, or global visual network, whereas fine-scale networks emerging at higher model orders reveal localized functional units such as V1/V2/V3 subdivisions, hand/foot/face motor regions, or distinct auditory subregions^15,19^. The single-scale approach limits the ability to capture the brain’s hierarchical and multi-scale functional architecture, which is essential for understanding the complexity of early brain development^16^. Emerging evidence indicates that functional brain organization arises from coordinated neural activity across multiple spatial scales, spanning local to global network levels^2,15,16,20–23^. Examining multiple scales in infancy is particularly critical, as different aspects of functional specialization and integration may manifest at different resolutions^2,15,20,23,24^. Finer-scale networks, for example, may be more pronounced in early infancy, associated with sensory and motor functions, whereas larger-scale networks, such as those related to higher-order cognitive processes, may mature more gradually^16^. A multi-scale approach therefore can provide complementary insights into early network maturation and offer a more complete framework for linking functional brain organization with developmental outcomes.

Collectively, these gaps highlight the need for more comprehensive and standardized multi-scale approaches^2,23,25^, to better capture the full spectrum of brain network development in early life. Standardization would facilitate cross-study comparisons and enhance the ability to conduct longitudinal research, ultimately enabling more accurate tracking of developmental trajectories. In recognition of this need, Iraji et al.^2^ developed the multi-scale NeuroMark 2.2 (NeuroMark_fMRI_2.2_modelorder-multi) template, a robust set of ICNs derived from rsfMRI data of over 100,000 individuals ranging from children to old adulthood, including both healthy and clinical populations^2^. Compared to its predecessor, this template provides improved granularity and generalizability, offering 105 canonical ICNs selected from an initial pool of 900 components obtained from a broad range of spatial scales through repeated ICA runs along with rigorous similarity and visual assessments^2,15,22^. These ICNs span both large-scale and fine-scale functional networks and are organized into seven functional domains and associated subdomains based on anatomical and functional characteristics^23^. By capturing functional organization across multiple spatial scales within a unified template, NeuroMark 2.2 provides a reproducible benchmark for cross-cohort and lifespan studies^2^. Despite its broad applicability, the systematic application of this multi-scale framework to infant populations remains largely unexplored, leaving it unclear whether and to what extent these canonical networks are present during early infancy.

To begin addressing these critical gaps, the present study provides a comprehensive investigation of the existence of multi-scale ICNs in infants aged 0–6 months using two complementary approaches for ICN identification. The burst independent component analysis (burstICA), a fully data-driven method, enables the unbiased discovery of infant-specific ICNs across spatial scales, without relying on predefined assumptions. This approach provides compelling evidence for the early emergence of complex, multi-scale network architecture. In parallel, we apply the NeuroMark framework, a semi-blind, template-guided method, to directly assess the presence of canonical adult ICNs in the infant data. By anchoring infant networks to a standardized multi-scale template, this approach facilitates robust cross-study and cross-age comparisons.

burstICA was developed to overcome key challenges in identifying functional networks during early development. It is a fully data-driven, model-order–agnostic approach designed to identify ICNs without dependence on spatial priors or predefined network templates. Unlike conventional ICA methods, which impose a fixed model-order constraint, burstICA performs iterative group-level ICA across a finely sampled continuum of decomposition dimensionalities (e.g., model orders ranging from 2 to 225 in increments of one), generating a large ensemble of decompositions spanning diverse spatial scales and hierarchical network configurations. Rather than treating each model order as an independent scale, burstICA leverages an ensemble-based selection strategy in which spatial scale is implicitly encoded by the originating model order of each ICN. This enables the simultaneous representation of both large-scale, spatially extended networks and fine-scale, localized subnetworks within a unified framework, while avoiding the redundant or unstable representations often associated with fixed-model-order analyses. To construct a robust set of ICNs, burstICA systematically evaluates all candidate components from the ensemble and selects, for each canonical ICN, the component exhibiting maximal spatial correspondence (highest spatial correlation) to a template (e.g., NeuroMark 2.2^2^). By integrating information across model orders through this scale-adaptive selection process, burstICA maintains the unbiased and fully blind nature of ICA while ensuring optimal matching across spatial resolutions. Importantly, burstICA extends the conceptual framework of multi-scale ICA (msICA) effectively capturing both fine-grained local networks and broadly distributed global architectures simultaneously^15^. This methodological flexibility is advantageous, given that different regions or systems may not conform to the same spatial scale within the brain’s functional hierarchy^2,15,22,26,27^, importantly in infant populations, where canonical networks are not yet fully established and different brain systems may mature at distinct spatial resolutions. Collectively, burstICA provides a rigorous and comprehensive analytic framework uniquely suited for mapping functional brain organization during infancy, supporting both exploratory discovery of novel networks and precise characterization of canonical ICNs. Detailed algorithmic descriptions and implementation procedures are provided in the Materials and Methods section.

Furthermore, the NeuroMark framework^28^ with the NeuroMark 2.2 functional template^2^ was applied to enhance the precision and interpretability of ICN identification. This semi-blind, template-informed approach uses spatially constrained ICA that facilitates refined delineation of infant ICNs while enabling the assessment of the presence or absence of specific canonical multi-scale networks at this early developmental stage. By aligning scan-specific ICNs with a standardized multi-scale reference, the NeuroMark framework supports direct and meaningful comparisons between the ICNs identified in infants and the canonical ICNs observed in older cohorts, enabling robust cross-study, cross-age, and cross-condition analyses.

The ICNs identified using burstICA and NeuroMark were then compared to the 105 canonical multi-scale ICNs identified in a large-scale, diverse population (N > 100,000) to assess whether these canonical networks are detectable in infants aged 0-6 months. To ensure validity and specificity, we conducted simulation-based analyses to confirm that the identified ICNs reflect the inherent structure of the data rather than methodological biases. burstICA was applied to the null dataset to assess the similarity of extracted components with the target ICN. Each canonical ICN from the NeuroMark 2.2 template served as a target ICN, against which the infant-derived and the null-derived components were evaluated. The spatial similarities between components from both the infant and the null datasets with target networks were quantitatively compared to evaluate statistical significance. Additionally, we tested whether the NeuroMark framework might enforce template-like components in the absence of true underlying signals by applying it to the null dataset. Furthermore, we assessed the specificity of individualized ICNs identified by the NeuroMark framework through a within-versus-between-network analysis, evaluating whether each network is significantly distinct from others for each scan. Lastly, to explore the functional architecture reflected by these ICNs, we calculated average static functional network connectivity (FNC) across the extracted ICNs. FNC, calculated as the Pearson correlation between ICN time courses, captures temporal relationships between networks, where strong positive or negative correlations indicate synchronous co-activation, respectively. Figure 1 provides a high-level overview of the analytical workflow for ICN identification and validation.

**Figure 1.**
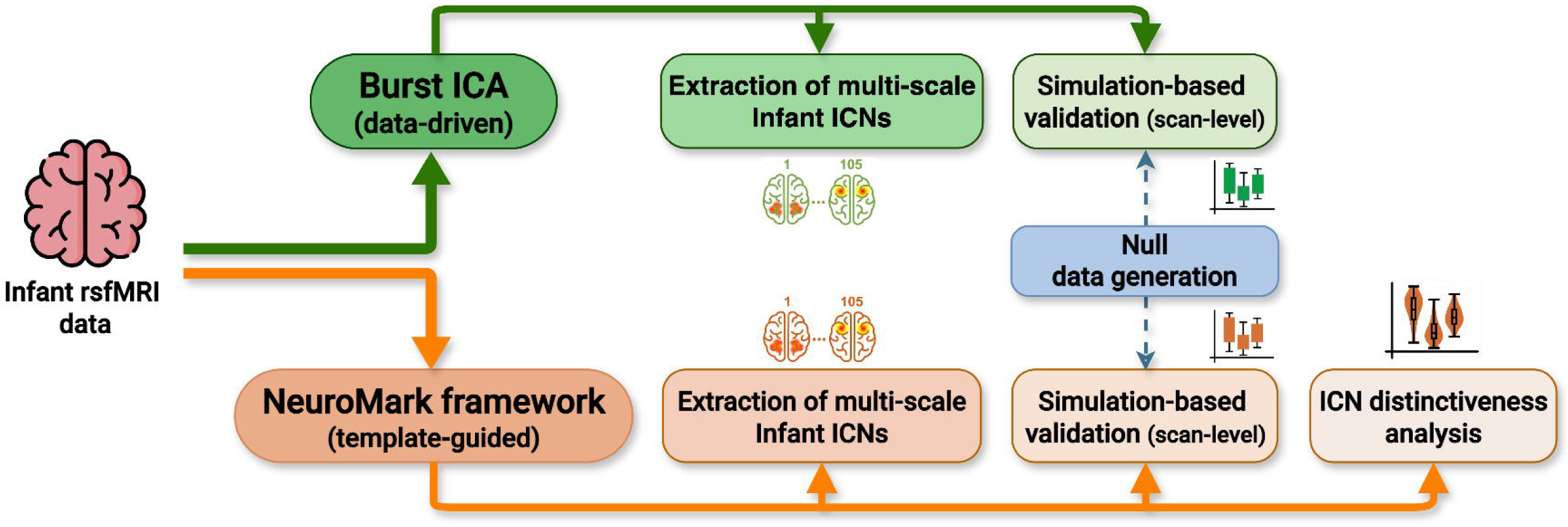
Overview of the Infant ICN Mapping and validation framework: This flowchart summarizes the overall analytic pipeline used to identify and validate multi-scale intrinsic connectivity networks (ICNs) in infant rsfMRI data. Two complementary approaches were applied: a fully blind, data-driven burstICA framework (top row, green) and the template-guided NeuroMark framework (bottom row, orange). Both methods were used to extract ICNs across multiple spatial scales. Simulation-based validation was performed using synthetic null data to assess the specificity of ICN estimates for each method. Additionally, NeuroMark-based ICNs were further assessed for their distinctiveness across domains.

The main contributions of this study include confirmation of the presence of multi-scale intrinsic connectivity networks (ICNs) in infants, the introduction of novel data-driven approaches that enable more reliable and accurate estimation of multi-scale ICNs, and validation of the NeuroMark framework for extracting multi-scale ICNs from infant resting-state fMRI data. The remainder of this paper is organized as follows. The Materials and Methods section describes the dataset, preprocessing procedures, burstICA implementation, and NeuroMark framework. The Results section presents ICN identification, validation, and connectivity findings. The Discussion section considers implications for early brain development and potential directions for future research

## Materials and Methods

### Participants

A total of 71 neurotypical infants (41 males, 30 females) were included from ongoing longitudinal studies conducted at the Marcus Autism Center in Atlanta, Georgia, USA (see Table 1 for demographic details). The mean gestational age at birth was 39.1 weeks (SD = 1.27). Infants were classified as neurotypical based on the absence of a family history of autism spectrum disorder in first-, second-, or third-degree relatives; no reported developmental delays in first-degree relatives; no pre- or perinatal complications; no history of seizures; and no known medical, genetic, auditory, or visual conditions. Infants with MRI contraindications were excluded. Written informed consent was obtained from a parent or legal guardian for all participants, and the study protocol was approved by the Emory University Institutional Review Board in accordance with all applicable ethical guidelines for research involving human participants.

**Table 1.**
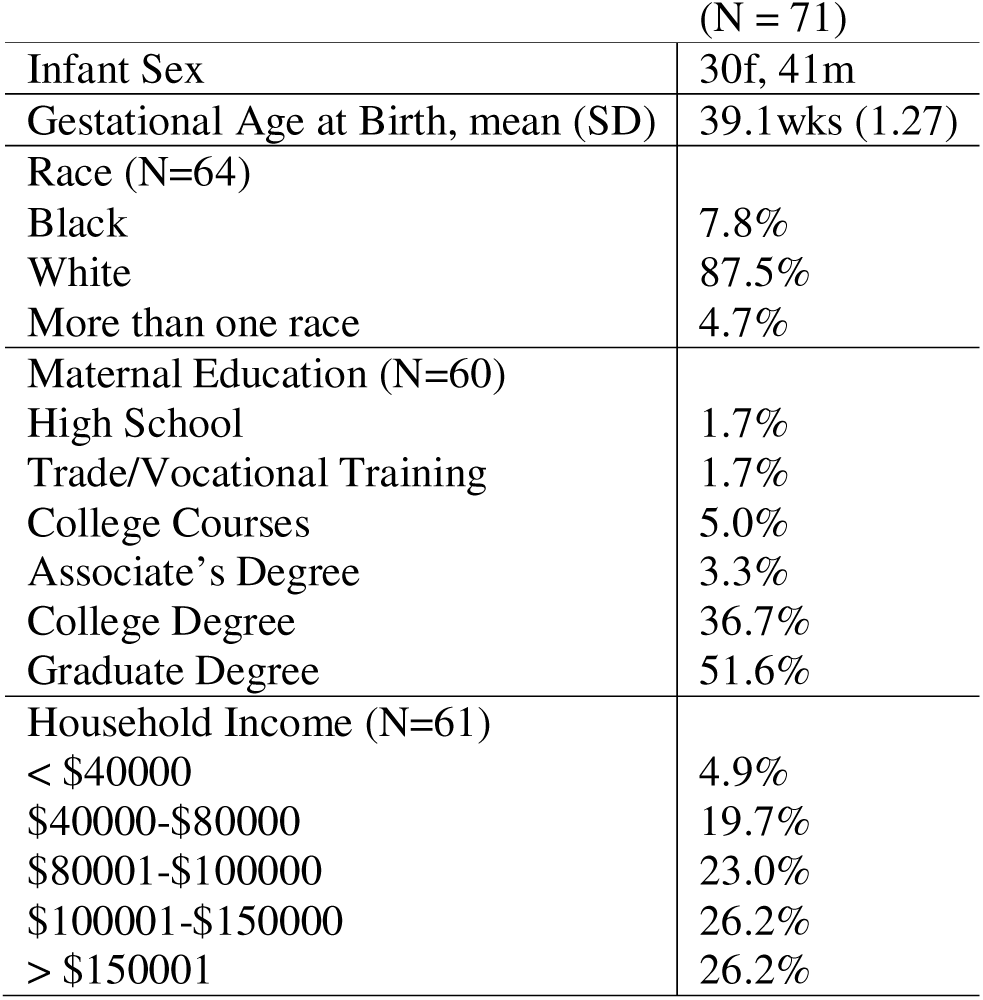
Demographics of participant sample. . For information categories with a participant number less than the total sample, the N is specified next to the category title.

### Data acquisition

rs-fMRI scans were acquired at Emory University’s Center for Systems Imaging Core using either a 3T Siemens Tim Trio or a 3T Siemens Prisma scanner, both equipped with a 32-channel head coil. Scans were scheduled at up to three pseudorandom time points between birth and nine months of age (in Supplementary Materials Fig. S1).

All imaging sessions were conducted during natural sleep. Infants were swaddled and soothed, typically through rocking or feeding, until asleep, then placed on a pediatric MRI bed. To minimize acoustic exposure (kept below 80 dBA), each infant wore sound-attenuating pediatric headphones with integrated MR-compatible optical microphones for real-time monitoring of in-ear noise levels. A custom-built acoustic hood was also placed inside the scanner bore. To mask scanner noise, white noise was gradually introduced through the headphones prior to scan onset. Throughout the session, infants were continuously monitored via an MRI-compatible camera (MRC Systems) mounted on the head coil. A trained researcher remained in the scanning room and paused the session if the infant awoke or if excessive noise was detected.

Multiband echo-planar imaging (EPI) sequences were used to acquire BOLD-sensitive rs-fMRI data. The scan parameters were Siemens Tim Trio (TR = 720 ms, TE = 33 ms, flip angle = 53°, field of view = 208 × 208 mm², 72 slices, 570 volumes (∼6 min 50 sec)) and Siemens Prisma (TR = 800 ms, TE = 37 ms, flip angle = 52°, field of view = 208 × 208 mm², 72 slices, 420 volumes (∼5 min 36 sec)). Each session included two rs-fMRI runs with anterior-to-posterior and posterior-to-anterior phase encoding directions to improve correction of susceptibility-induced distortions.

### Preprocessing

We performed standard fMRI preprocessing using the FMRIB Software Library (FSL v6.0, https://fsl.fmrib.ox.ac.uk/fsl/fslwiki/) and the Statistical Parametric Mapping (SPM12, http://www.fil.ion.ucl.ac.uk/spm/) toolboxes under the MATLAB 2020b environment. Firstly, the initial ten dummy scans with substantial signal changes were discarded. Following that, we corrected the distortion in the images using SBRef data with phase encoding blips in reverse. After distortion correction, we performed slice timing to correct for these slice-dependent delays. Head motion correction was performed followed by the slice timing to realign all the scans to the reference scan. To normalize the infant data to the standard adult Montreal Neurological Institute (MNI) space, we applied a two-step normalization procedure. We first warped the UNC-BCP 4D Infant Brain Template (https://www.nitrc.org/projects/uncbcp_4d_atlas/)^29^ into the adult MNI space using the EPI template as the reference. Here, to mitigate the introduction of an age-specific bias into normalization, we chose the template with the age of 4 months, which is the median of the age range of our infant dataset. After having the UNC-BCP template in the adult MNI space, we normalized the infant data using it as the reference. Finally, the normalized fMRI data were spatially smoothed using a Gaussian kernel featuring a 6 mm full width at half maximum (FWHM).

### Multi-scale ICN Estimation and Validation

#### Burst Independent Component Analysis

burstICA framework consists of three sequential stages: (i) multi-model-order ICA decomposition and ensemble generation, (ii) component selection via template matching, and (iii) scan-level reconstruction and FNC estimation. Figure **2** demonstrates the overview of the burstICA framework.

**Figure 2.**
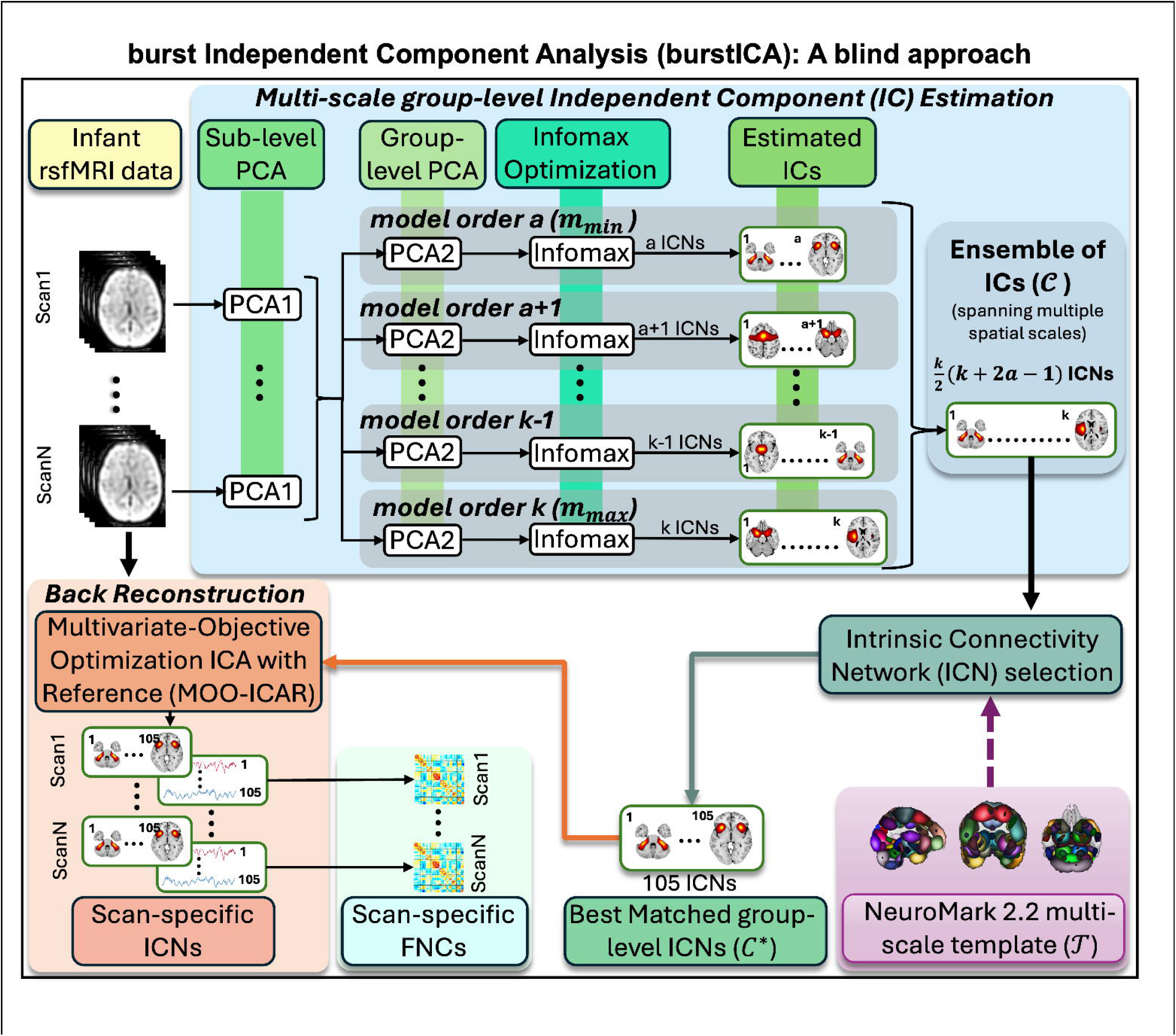
Overview of the burst independent component analysis (burstICA) framework. Group-level data are iteratively decomposed using PCA and Infomax ICA across a finely sampled range of model orders, yielding an ensemble of independent components *(ICs)* spanning multiple spatial scales. For each canonical ICN in the NeuroMark 2.2 template, the component exhibiting the highest spatial correlation is selected, resulting in a robust set of representative group-level ICNs. These ICNs are subsequently used as spatial references for s*can*-level reconstruction using multi-objective optimization ICA with reference (MOO-ICAR), producing scan-specific spatial maps and time courses. Functional network connectivity (FNC) is then estimated from the reconstructed time courses using pairwise correlations.

#### ICA decomposition and Ensemble Generation

Given the preprocessed and temporally concatenated group-level rsfMRI data *X ∈ R^VxT^*, an ICA decomposition is repeatedly applied across a finely sampled range of model orders *m*. At each model order, dimensionality reduction was performed using subject- and group-level principal component analysis (PCA), followed by group-level ICA using the Infomax algorithm (see Supplementary Methods).

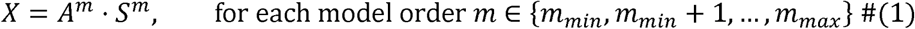

Where *A^m^ ∈ R^Txm^*, denotes the associated IC time courses, and *S^m^ ∈ R^mxV^* denotes the spatial maps (mixing matrix) at model order *m*.

This procedure yielded a large ensemble of ICs spanning multiple spatial scales and network configurations:

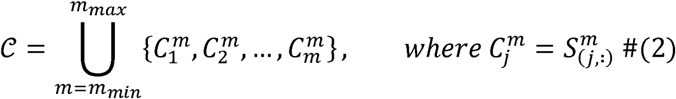

#### Component Selection via Template Matching

Given the canonical ICN template *T = {T_1_,T_2_,…,T_K_}* (e.g., NeuroMark 2.2), we performed a Specifically, for each canonical ICN T_k_, the IC from the ensemble c with the highest spatial spatial matching procedure to select the best representative IC for each canonical network. correlation to the template *T_k_* is selected:

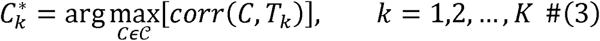

The final set of robustly identified ICNs thus becomes:

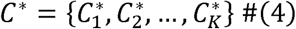

This ensemble-based strategy ensures blind, scale-adaptive component selection while maintaining high spatial correspondence to biologically interpretable networks.

#### Scan-Level Reconstruction and FNC Estimation

Individual-level ICNs were reconstructed using multi-objective optimization ICA with reference (MOO-ICAR)^28^, a spatially constrained ICA approach that uses group-level ICNs as soft priors. This method preserves subject-level variability while aligning individual maps to population-level components. The corresponding time courses were used to estimate static FNC, defined as pairwise Pearson correlations between ICN time series.

#### NeuroMark framework: A Reference-Informed Approach

We estimated ICNs through a reference-informed ICA, NeuroMark framework, employing MOO-ICAR algorithm. The NeuroMark 2.2 multi-scale template^2^, publicly available from the TReNDS Center website (https://trendscenter.org/data/), served as the spatial reference to guide the identification of ICNs in our infant resting-state fMRI dataset. MOO-ICAR^28^ was conducted using the GIFT toolbox (https://github.com/trendscenter/gift)^30^. This approach enables scan-specific estimation of ICNs while maintaining correspondence across scans.

For a given scan *k*, the preprocessed rsfMRI data matrix is denoted as *X^k^ ∈ R^VxT^*, where *V* is the number of voxels and *T* is the number of time points. For each template ICN *S_1_ ∈ R^VxT^*, the corresponding scan-specific component *S^k^_l_* is estimated as:

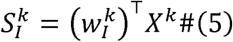

where *W^k^_l_ ∈ R^VxT^* is the unmixing vector to be optimized. MOO-ICAR^28^ simultaneously optimizes the following two objectives:

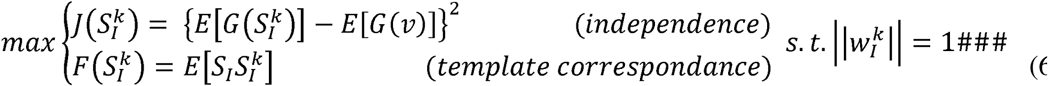

Here, *v ∽ N(0,1)* is a Gaussian variable, *G(.)* is a non-quadratic function, and *E(.)* denotes expectation. The final cost function is the weighted sum of both objectives:

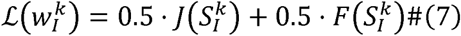

Solving this optimization yields one scan-specific ICN for each template network, resulting in *N* scan-specific ICNs per scan. This procedure preserves scan-specific variability while Solving this optimization yields one scan-specific ICN for each template network, resulting ensuring correspondence with standardized template networks.

Spatial correspondence between estimated ICNs and template ICNs was quantified using Pearson correlation. Network specificity was assessed using within-versus-between-network similarity comparisons. The associated time courses were subsequently used to compute static FNC, capturing inter-network temporal dependencies. Figure 3 illustrates the pipeline figure of the NeuroMark framework.

**Figure 3.**
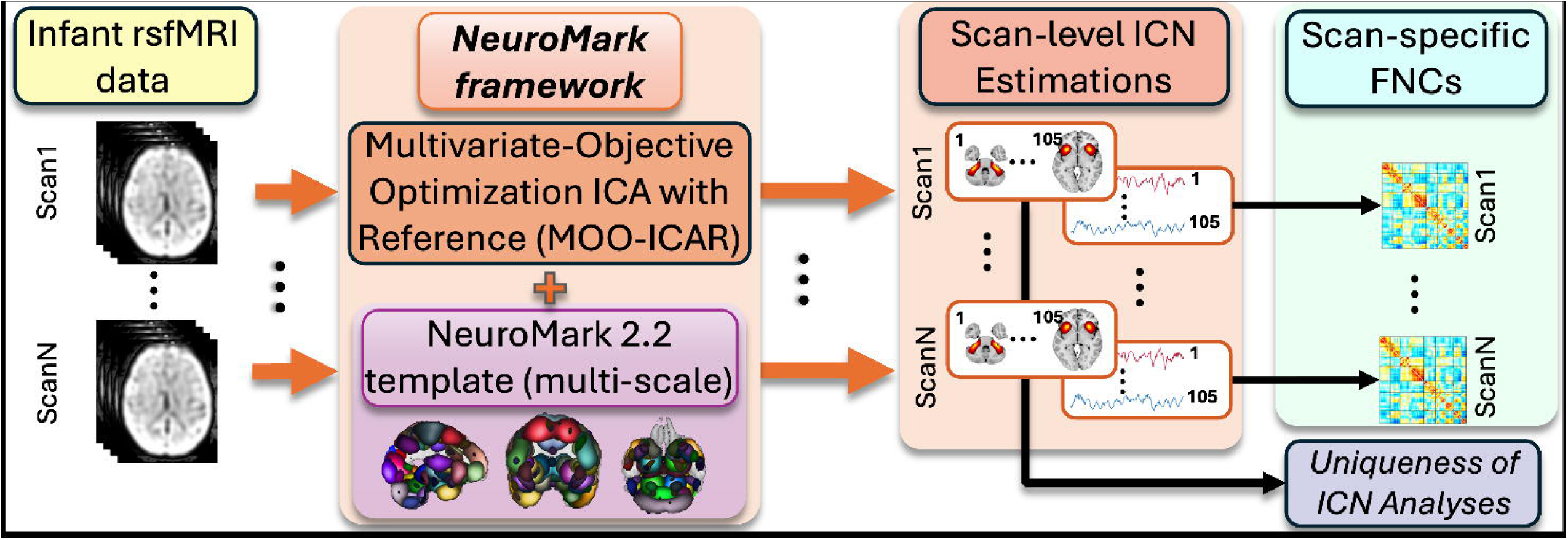
Overview of NeuroMark framework for reference-informed scan-level ICN estimation. Integrates MOO-ICAR method and the NeuroMark 2.2 template as a reference to generate precisely 105 scan-level ICN estimations from the infant dataset. The spatial maps are further used to analyze the unique information in each ICN, and the time courses are used to evaluate FNC.

## Results

As shown Figure 4, group-level burstICA (left) and the NeuroMark framework (right) produced coherent domain-wise groupings of spatial maps, revealing reproducible network architecture across the infant cohort.

**Figure 4.**
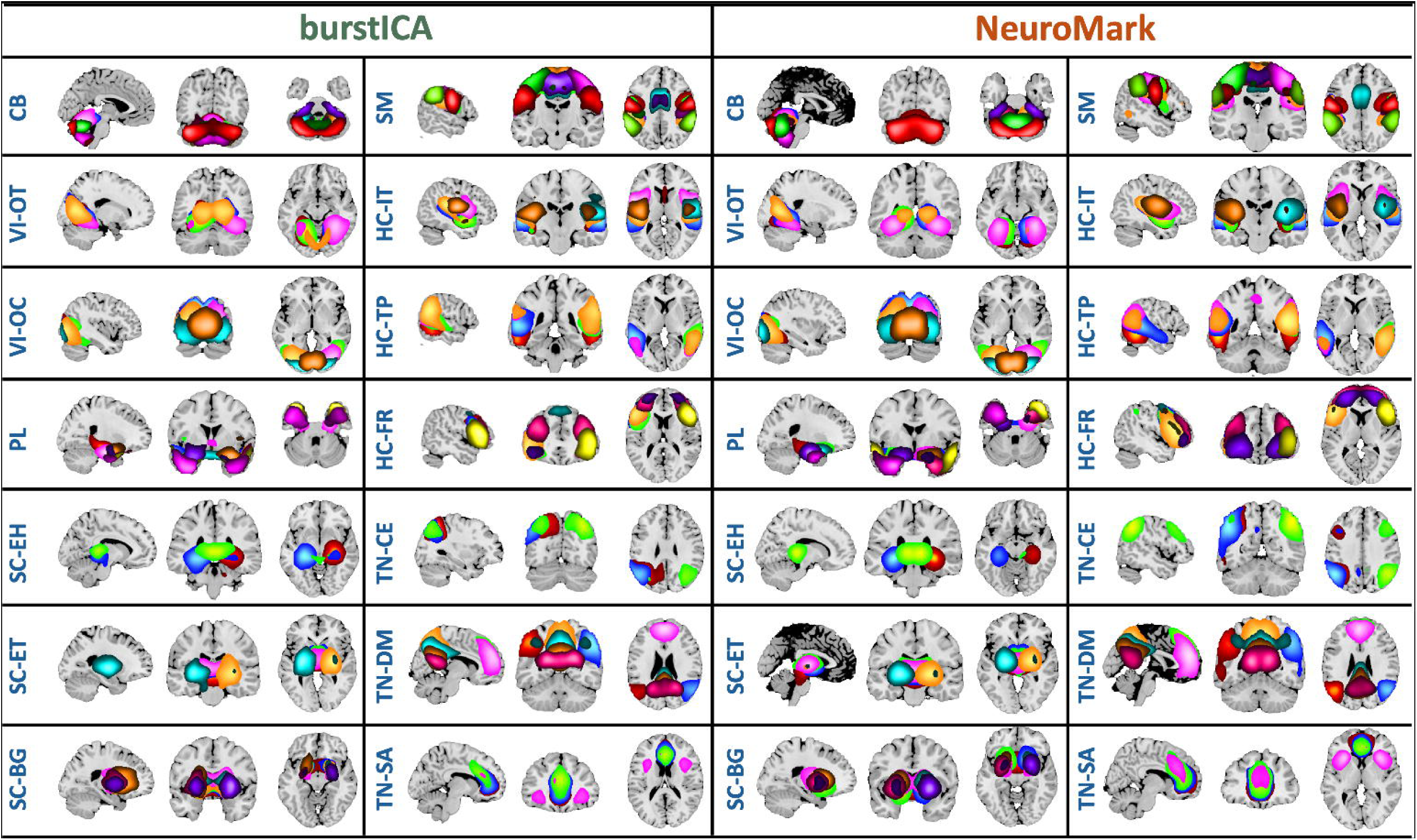
Multi-scale intrinsic connectivity networks (ICNs) identified from infant resting-state fMRI. Shown are 105 ICNs estimated using two complementary approaches: best matched group-level burstICA (left) and average ICNs from the NeuroMark framework. Networks are organized into 7 functional domains and 14 subdomains: Cerebellar (CB); Visual (VI-OT, occipitotemporal; VI-OC, occipital); Paralimbic (PL); Subcortical (SC-EH, extended hippocampal; SC-ET, extended thalamic; SC-BG, basal ganglia); Sensorimotor (SM); Higher Cognition (HC-IT, insular temporal; HC-TP, temporoparietal; HC-FR, frontal); and Triple Network (TN-CE, central executive; TN-DM, default mode; TN-SA, salience).

### Blind msICA-based Identification of Infant ICNs via burstICA

#### Extraction of multi-scale Infant ICNs: spatial correspondence and FNC similarity with canonical “adult-like” networks

The spatial similarity between each infant-derived ICN using burstICA and its corresponding template ICN was measured using Pearson correlation, with all correlations exceeding rho > 0.5. Notably, four ICNs exhibited exceptionally high spatial similarity to their template counterparts, with correlation coefficients (rho) > 0.9, three of these belonged to the sensorimotor (SM) domain and one to the high cognitive–insular temporal (HC–IT) domain. In total, 32 of the 105 ICNs had spatial correlations exceeding rho > 0.8, while the remaining 73 ICNs all exceeded rho > 0.5. The full range of spatial correlation coefficients spanned from 0.5456 to 0.9587, with an average of 0.7462 ± 0.0962, surpassing the matching threshold used in previous studies^28,31^. Figure 5a summarizes these spatial similarity scores, while Figure 5b shows the distribution across ICNs. The FNC matrix Figure 5c, revealed a broad spectrum of inter-network interactions, with connection strengths ranging from –0.48 to 0.99. The overall average within-subdomain FNC was 0.5082, while the average between-subdomain FNC was –0.0137. Notably, comparison of the group-average FNC matrix from burstICA with that of a large-scale adult cohort (UK Biobank^23^, N = 39,342) yielded a correlation of 0.6917, indicating a substantial preservation of modular network structure from infancy through adulthood.

**Figure 5.**
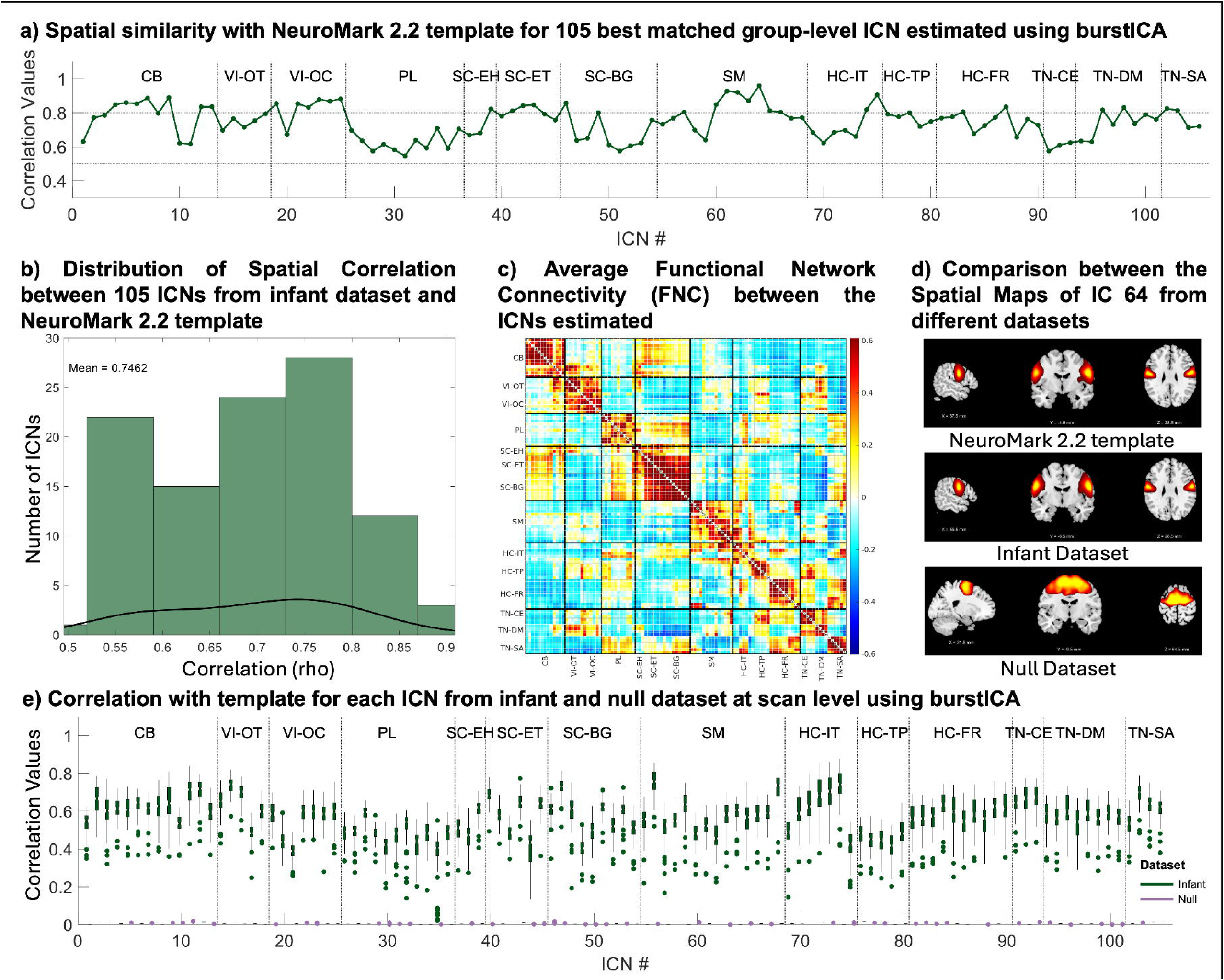
Analysis of 105 ICNs estimated from the infant dataset from burstICA and matched with the NeuroMark 2.2 Template. **a)** Spatial similarity using Pearson correlation (rho values) for each of the 105 best-matched group-level ICNs estimated using burstICA, compared with the NeuroMark 2.2 template. This panel illustrates the individual correlation values indicating how well each ICN aligns with the template b) Distribution of spatial correlation (rho values) with the template for the 105 ICNs from the infant dataset. This histogram shows the range and frequency of similarity values, providing an overview of the alignment between the estimated ICNs and the template. c) Average FNC between the ICNs derived from the infant dataset. This panel presents the mean connectivity patterns among the identified ICNs, offering insights into their functional relationships. d) Comparison of spatial maps for independent component (IC) 64 from different datasets: NeuroMark 2.2 template, the infant dataset, and a null dataset. This example highlights the similarities and differences in ICNs obtained from the actual data versus the null dataset. e) Scan-level ICNs reconstructed using the 105 best-matched ICNs. This panel shows the correlation with the template for scan-level ICNs obtained using burstICA from both the infant and null datasets, demonstrating the reliability of the ICNs in the infant dataset.

#### Simulation-Based Validation of Multi-Scale ICNs Detected using burstICA in Infant Data

Following the estimation of ICNs using burstICA, we aimed to evaluate the specificity of the identified infant networks and determine whether they reflected meaningful brain organization rather than arising by chance. To this end, we compared spatial correlations between the NeuroMark 2.2 template, and the 105 best-matched components derived from both the infant dataset and a null dataset generated using the same burstICA pipeline. The null dataset was created through a simulation framework that preserved the structural properties of fMRI data while excluding ICNs of interest. Low-correlation spatial maps were paired with synthetic time courses to generate pseudo-scans, ensuring that no meaningful ICN structure was embedded (see Supplementary Materials for details). A substantial drop in correlation within the null data would indicate that the infant-derived ICNs reflect meaningful, non-random brain structure.

The average spatial correlation between the target ICNs and their best-matched components from the null data was 0.0056 ± 1.56×10⁻, compared to 0.5710 ± 0.0140 in the actual infant data (Figure 5e). This stark contrast underscores the presence of meaningful, biologically relevant structure in the infant-derived ICNs. Paired t-tests yielded t-values ranging from 26.46 to 124.54 and p-values < 3.42×10⁻³, indicating that the ICNs identified in the infant rsfMRI data are significantly more aligned with the canonical NeuroMark networks than components arising from null distributions. Importantly, the correlations shown in Figure 5e reflect scan-level ICN estimates, where each box plot represents the distribution of correlations for a given ICN across individual scans. This visualization highlights the expected scan-to-scan variability in spatial similarity, particularly given that the scans were acquired from infants spanning a broad developmental window (0–6 months). As expected, such variability is absent in the null dataset, which exhibits uniformly negligible correlations. Figure 5d illustrates this comparison for a representative ICN (ICN 64), where the spatial map from the infant data shows significantly higher similarity to the corresponding template ICN than the matched component from the null data. In both datasets, the matched component refers to the one exhibiting the highest spatial correlation with the target ICN from the NeuroMark 2.2 template. Together, these findings strongly support the specificity and biological validity of the ICNs estimated to be from early infancy, considering the fully data-driven nature of burstICA.

### Reference-Informed Identification of Adult-like ICNs in Infants: NeuroMark framework

#### Extraction of multi-scale Infant ICNs: spatial correspondence and FNC similarity with canonical “adult-like” networks

The multi-scale ICNs estimated using NeuroMark framework showed a significant boost in spatial correspondence with the template (rho > 0.83; mean ± SD: 0.9138 ± 0.0236), with the highest reaching rho = 0.96 (Figure 6a and Figure 6b). These results demonstrate robust replication of canonical networks in the infant brain, supporting the framework’s superiority and applicability in future infant studies.

**Figure 6.**
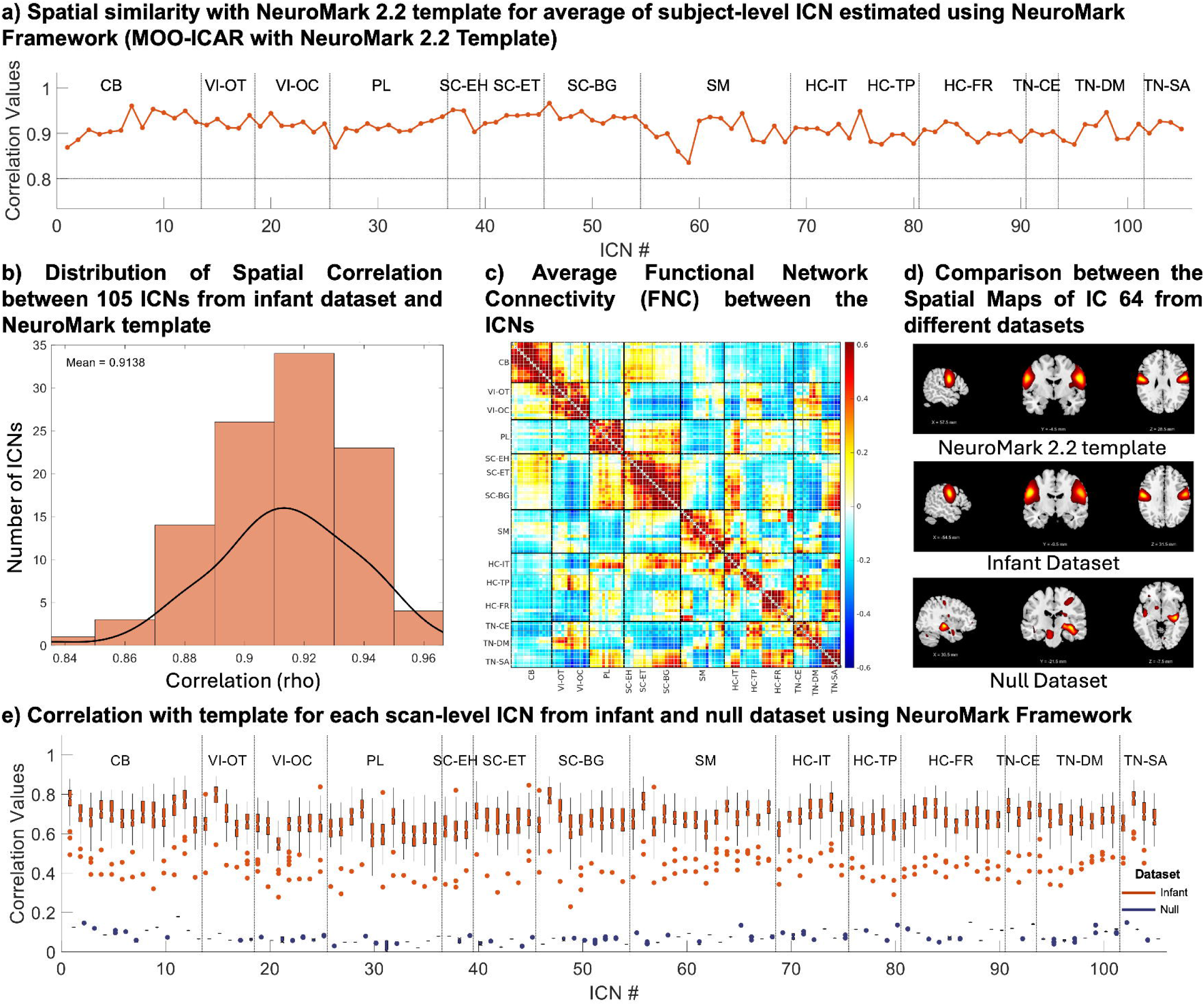
Analysis of 105 ICNs Estimated Using the NeuroMark framework from the Infant Dataset. **a)** Spatial similarity between the average estimated scan-level ICNs and the NeuroMark 2.2 template. This panel shows the mean correlation values for ICNs across scans, highlighting their alignment with the template. b) Distribution of spatial correlation between the 105 ICNs from the infant dataset and the template. This histogram illustrates the range and frequency of similarity values, providing an overview of how well the estimated ICNs match the template. c) Average FNC among the ICNs estimated using NeuroMark framework. This panel presents the mean connectivity patterns between the identified ICNs, offering insights into their functional relationships d) Comparison of spatial maps for IC 64 across different datasets: NeuroMark 2.2 template, the infant dataset, and the null dataset. e) Correlation with the template for each scan-level ICN obtained using NeuroMark framework from both the infant and null datasets.

Average FNC across the scans revealed connection strengths widely ranging from 0.48 to 0.96 (Figure 6c). The overall mean within-subdomain FNC was 0.5244, and the mean between-subdomain FNC was 0.0155. Further comparison with the adult UK Biobank FNC matrix yielded a substantial correlation of 0.7166, reinforcing the high degree of modular similarity between infant and adult network architecture.

#### Simulation-Based Validation of Multi-Scale ICN Detected using Neuromark framework in Infant Data

To further assess the NeuroMark framework, we evaluated whether the method could potentially enforce the generation of template-like ICNs even in the absence of true signal. Specifically, we applied the NeuroMark pipeline to the same null dataset used in the burstICA specificity analysis. This allowed us to determine whether the framework might extract components that spuriously match the template due to algorithmic bias, rather than reflecting underlying neural organization.

As expected, the spatial similarity between the ICNs estimated from the null dataset and the NeuroMark 2.2 template was substantially lower than that observed in the infant data. Across all networks, ICNs derived from the infant data exhibited strong spatial correspondence with the template, while the null-derived component showed minimal similarity. Figure 6e presents Pearson correlation coefficients between each template ICN and its best-matched component from both the infant and null datasets at the scan level. Spatial similarity for the infant data was significantly higher (mean ± SD: 0.6737 ± 0.0082) compared to the null data (mean ± SD: 0.0781 ± 0.0311). Paired t-tests for each ICN revealed highly significant differences, with t-values = 40.54 to 88.41 and all p-values < 7.11 × 10⁻lJ¹, confirming the robustness and specificity of the ICNs estimated from the infant data. As in the burstICA analysis, the correlations shown in Figure 6e represent scan-level ICN estimates, with each box plot illustrating how spatial similarity varies across scans for a given ICN. Figure 6d illustrates an example ICN (ICN 64), demonstrating the difference in the spatial maps derived from the NeuroMark 2.2 template, the infant dataset, and the null dataset. The infant-derived ICN closely matches the template, while the null-derived component does not. Notably, strong spatial correspondence was observed even in scans acquired during the first week after birth (mean correlation = 0.6286 ± 0.0478), supporting the early emergence of canonical functional network architecture in infancy. These findings confirm both the accuracy of ICN estimation in infants and the practical utility of the NeuroMark framework in infant rsfMRI data.

#### ICN distinctiveness analysis: Scan-level network specificity using within-versus between network comparison in Infants

Building on our null-model analysis, which demonstrated that spatial correlations between null component maps and the NeuroMark 2.2 templates are highly distinct from the levels observed for true component matches and thus ruled out chance alignment, we next assessed whether each infant scan retained the unique spatial signature of its matched ICN, despite the presence of other, similar networks in the template. For every target ICN (X), we first identified its closest non-matched peer termed the “comparison ICN” (Y), as the ICN exhibiting the second-highest spatial overlap with X in the template. For each scan, we extracted both the matched ICN (X) and its comparison ICN (Y) and calculated their respective Pearson correlations with the target template ICN (X). This matched-versus-comparison analysis constitutes a within-(matched) versus between-network (comparison) evaluation at the scan level.

Across all ICNs and scans, the matched ICN consistently exhibited higher spatial similarity to the template than the comparison ICN. As illustrated in Figure 7-top, the correlation difference was consistently positive across all ICNs (mean ± SD = 0.2618 ± 0.1288), indicating robust identification at the individual-scan level. Paired t-tests further confirmed these differences as statistically significant for all ICNs (t-values: 15.04 to 173.43, p < 1.54 × 10⁻lJlJ, Figure 7-bottom).

**Figure 7.**
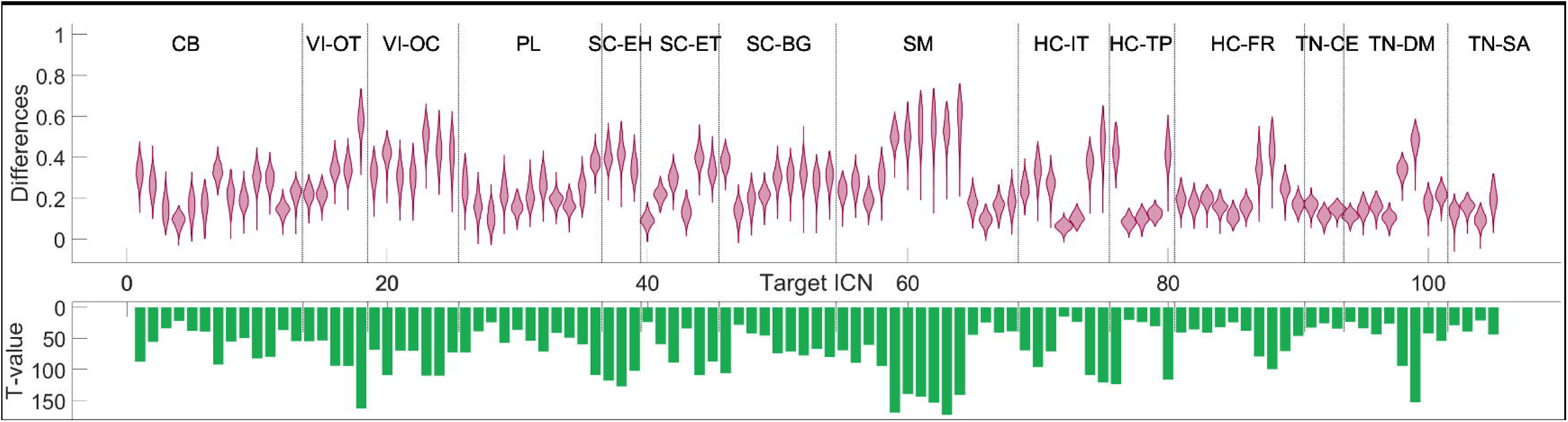
Distinctiveness of scan-level ICNs relative to template networks. The top panel shows the distribution of differences in spatial similarity between each estimated scan-level network of interest and its matched template ICN, compared to similarity with the most similar non-matched (comparison) ICN. The bottom panel displays the corresponding t-values from paired t-tests, demonstrating that matched networks show significantly higher similarity to the template than comparison ICNs.

These findings highlight that, even within a multi-scale template comprising inherently similar networks, the ICNs matched for each infant scan consistently provided the most precise and reproducible spatial representations. Together with earlier group-level findings, this scan-level analysis confirms that ICNs were not only reliably detected across infants but also accurately matched on a per-scan basis, thus supporting the reliability of the method.

## Discussion

While rsfMRI studies in adults have provided robust characterizations of ICNs, analogous efforts in infancy remain limited. Prior infant studies have identified several ICNs^7–9,11,12,32–34^, yet only a subset have been thoroughly examined^6–9^, and no standardized, multilJscale networks have yet been established. This gap has constrained robust crosslJstudy and crosslJage comparisons. Here, we bridge that divide by mapping 105 canonical multilJscale ICNs in infants using the NeuroMark 2.2 template^2,23^. Because NeuroMark 2.2 is used in adult work, our infant maps can be directly aligned with adult ICNs, enabling immediate assessment of which networks are truly “adultlJlike” in early life and facilitating longitudinal, lifespan analyses.

In this study, we employed two complementary approaches, burstICA and the NeuroMark framework, to generate a comprehensive, multi-scale map of infant ICNs. First, to determine whether canonical ICNs could be extracted purely from infant data itself, we applied burstICA, a blind msICA-based pipeline designed to maximize sensitivity. This data-driven approach resulted in spatial correlations exceeding rho > 0.5 across all 105 ICNs surpassing the previously reported thresholds^28,31^. These results provide compelling evidence that adult-like network topologies are already present by six months of age. Importantly, simulation-based validation using null datasets confirmed that the burstICA-derived ICNs were biologically specific rather than methodological biases. Subsequently, we aimed to test if presenting a predefined template through reference-informed approach contribute for cross-age study. Thus, we applied the reference-informed NeuroMark framework, to produce precise, template-guided infant ICNs. We observed even higher spatial correspondence (rho > 0.8) using a standardized template-guided approach, and rigorous specificity testing (including null data and within-versus between-network comparisons) confirmed that each identified ICN retained a distinct and reproducible spatial profile, highlighting the reliability of the NeuroMark-informed ICNs.

Our findings offer critical insights into neurodevelopment, particularly the early formation of brain network architecture. Importantly, the high spatial correlations observed in our infant sample are comparable to those reported in independent adult cohorts in Iraji et al.^2^, where ICN-to-template correlations ranged from 0.80 to 0.95. This striking spatial similarity indicates that the macroscale topography of functional networks is largely established early in life, and that subsequent developmental processes during infancy likely involve refinement of functional specialization and increases in connectivity strength rather than substantial alterations in network spatial organization. While prior research demonstrated complex networks, such as the DMN emerge in a rudimentary form and mature over the first year, with the salience network developing by the second year^11,35^, our results indicate that ICNs including the DMN and salience network in infants aged 0 to 6 months already exhibit a high degree of similarity to their mature counterparts. Specifically, the multi-scale ICNs we identified showed that DMN-related networks within the TN-DM subdomain have a high spatial similarity to the template (mean rho = 0.9047 ± 0.0245). Similarly, the TN-SN subdomain related to the salience network showed substantial similarity (mean rho = 0.9149 ± 0.0123). These correlations were comparable to those observed for primary networks, such as visual areas, which are expected to show adult-like structure even in neonates^11,35^, VI-OT (mean rho = 0.9226 ± 0.0124) and VI-OC (mean rho = 0.9202 ± 0.0126). Collectively, these findings suggest that the anatomical foundations or structural frameworks underlying these networks are largely established in early infancy. Subsequent developmental refinement likely involves fine-tuning functional connectivity and enhancing behavioral integration, as supported by Gao et al. (2015), Emerson et al. (2016), and other longitudinal studies^10,35^. Additionally, our use of multi-scale decomposition likely enhanced the sensitivity to detect these patterns, which may otherwise appear more diffuse when assessed using traditional single-scale approaches.

We also observed exceptionally high similarity within subcortical ICNs, SC-BG (mean rho = 0.9380 ± 0.0128), SC-EH (mean rho = 0.9348 ± 0.0274), and SC-ET (mean rho = 0.9346 ± 0.0089). These values suggest a high degree of spatial fidelity in subcortical networks during early infancy. Previous studies have successfully delineated subcortical connectivity, including the basal ganglia^36,37^, the thalamus^38,39^, and the hippocampus^40,41^—confirming that these regions are functionally active from birth and undergo substantial reorganization. However, the spatial patterns reported in these studies often appeared in a more primitive or less differentiated form. Our findings extend this body of work by demonstrating, for the first time, that when examined through a multi-scale framework aligned with a high-resolution template derived from older participants, subcortical ICNs in infants not only emerge consistently but do so with spatial precision comparable to that observed in mature brain networks. This may reflect both the early functional maturation of these deep structures and the enhanced sensitivity of multi-scale decomposition to capture their architecture.

Another key finding concerns the global organization of the infant whole-brain functional connectome, as captured by our FNC analyses. Infant FNC patterns derived from both burstICA and NeuroMark analyses demonstrated substantial similarity to adult FNC profiles from the UK Biobank dataset (burstICA: rho = 0.6917; NeuroMark: rho = 0.7166), indicating that core features of adult-like, system-level functional architecture, including long-range interactions across distributed cortical systems, are already emerging within the first six months of life. Notably, the NeuroMark framework consistently yielded stronger spatial and FNC correspondence compared to burstICA, highlighting the advantages of template-guided ICN approaches for estimating corresponding functional patterns and enhancing reliability and cross-dataset comparability. Within-domain connectivity was robust and positive across both methods (burstICA: mean FNC = 0.5082; NeuroMark: 0.5244), closely mirroring adult within-domain values (UKB: 0.4186). Furthermore, clear anti-correlated functional connectivity blocks emerge between primary sensory systems (e.g., SC, VI, and SM) and higher-order transmodal networks, such as the salience (TN-SA), cognitive control (TN-CE), and default mode (TN-DM) domains. These long-range, cross-system antagonistic interactions reflect an early instantiation of functional segregation and hierarchical organization, key hallmarks of the mature connectome^42,43^. Consistent with prior developmental studies, we observe early manifestations of network-level segregation and integration, particularly evident within sensory-motor and proto-default mode regions^11,32,44,45^. Although these signals are typically weak and less differentiated in infancy, they nonetheless signal the foundational principles of connectome maturation that strengthen markedly with age.

Additionally, our approach overcomes a long-standing limitation in ICA-based analyses: the challenge of model order selection. Traditional methods often rely on fixed or heuristic model orders, which may restrict the resolution or completeness of network detection. In contrast, burstICA systematically explores a finely grained range of model orders, enabling more comprehensive and unbiased identification of ICNs. While this study focused on ICNs that aligned closely with NeuroMark 2.2, burstICA offers future potential for discovering multi-scale networks unique to infancy beyond the scope of this paper. In parallel, we demonstrated the feasibility of applying the NeuroMark framework and the generalizability of the NeuroMark 2.2 template to infant data. The computational burden of burstICA can be substantial for large-scale datasets, while the template-guided NeuroMark approach is more efficient when a robust reference is available. Despite these trade-offs, both methods reliably identified ICNs in infant data, underscoring their potential for advancing early neurodevelopmental assessment. These tools may be particularly valuable for identifying early deviations in brain network development in infants at elevated likelihood for neurodevelopmental conditions such as ASD or ADHD, providing a foundation for characterizing divergent neurodevelopmental trajectories during infancy.

These findings collectively emphasize the need to study infant brain development with greater resolution and scale. They also lay groundwork for future efforts to evaluate how network organization varies with age and to prioritize longitudinal studies that track developmental trajectories from infancy through early childhood, enabling a more complete characterization functional network maturation. Scaling larger and more diverse samples will be critical for enhancing generalizability. In addition, applying these approaches in clinical or high-risk populations may facilitate earlier identification of neurodevelopmental vulnerabilities and opportunities for intervention. Finally, integrating dynamic FNC analyses could provide deeper insight into the evolving organization of the developing brain.

## Conclusion

In this study, we provide the first comprehensive, multi-scale characterization of ICNs in early infancy that is directly comparable to mature functional brain organization. By integrating a fully data-driven burstICA framework with the reference-informed NeuroMark approach, we demonstrate that a broad repertoire of canonical ICNs, spanning cortical, subcortical, and transmodal systems, can be robustly identified in infants as young as 0–6 months. Across 105 networks, infant ICNs exhibited striking spatial correspondence with the NeuroMark 2.2 template, indicating that the macroscale topology of the functional connectome is largely established within the first half-year after birth.

Our findings challenge the prevailing view that higher-order association networks emerge only gradually over the first years of development. Instead, we show that networks associated with the default mode, salience, and cognitive control systems already display adult-like spatial organization in early infancy, with similarity comparable to that observed in matured cohorts. Subcortical ICNs exhibited particularly high spatial fidelity, highlighting the early functional maturation of deep brain structures and underscoring the value of multi-scale decomposition for resolving their architecture. At the systems level, infant functional network connectivity patterns revealed clear signatures of within-domain coherence, long-range interactions, and emerging anti-correlations between primary sensory and transmodal networks, hallmarks of hierarchical organization that foreshadow the mature connectome.

Methodologically, this work addresses key limitations of prior infant rsfMRI studies by overcoming model-order constraints and enabling standardized, cross-age comparisons. BurstICA provides a sensitive, unbiased means of discovering multi-scale networks directly from infant data, while NeuroMark offers a computationally efficient and highly reproducible framework for aligning infant and adult ICNs within a common reference space. Together, these complementary approaches establish a scalable foundation for longitudinal and lifespan investigations of functional brain development.

By defining a reliable, template-aligned atlas of infant ICNs, this study lays critical groundwork for future research into developmental trajectories and early neurobiological vulnerability. Applying these methods to longitudinal cohorts and high-likelihood clinical populations may enable earlier detection of atypical network development and inform the timing of intervention. More broadly, our results suggest that postnatal brain maturation is characterized less by the emergence of entirely new networks and more by the progressive refinement and strengthening of a functional architecture that is already remarkably adult-like in early infancy.

## Data availability

Due to IRB restrictions, the raw infant MRI data analyzed in this study cannot be publicly shared. Access to these data may be granted upon reasonable request and subject to Institutional Review Board approval; requests should be directed to Dr. Sarah Shultz.

## Supplementary material

Supplementary material is available at *Brain* online

