## Supplementary material for "Born Connected: The Early Emergence of Adult-Like Multi-Scale Brain Networks in Infancy"

**The PDF file includes:**

- Materials and Methods
    - Dataset
    - Preprocessing
    - Estimating ICN from rsfMRI
        - burst Independent Component Analysis (burstICA)
        - NeuroMark framework: A Reference-Informed Approach
    - Functional Network Connectivity
- Fig S1 to S3
- Table S1

### Materials and Methods

Dataset

Participants were typically developing infants enrolled in prospective longitudinal studies of infants at low- and elevated familial genetic likelihood for autism spectrum disorder, conducted at the Marcus Autism Center in Atlanta, Georgia, USA. Typicality was ascertained by the absence of factors associated with increased likelihood of developmental disability: all infants had uncomplicated deliveries (mean gestational age = 39.1 weeks, SD=1.27 weeks), had no family history of ASD in first, second, or third degree relatives, no developmental delays in first degree relatives, no pre- or perinatal complications, no history of seizures, no known medical conditions or genetic disorders, and no hearing loss or visual impairment. Participants had no contraindications for MRI. Prior to participation, informed written consent was acquired from the parents of all participants, and ethical approval for all study procedures was granted by the Emory University Institutional Review Board. The rsfMRI data was acquired using a 32-channel head coil, with scans conducted during natural sleep. Data was collected using a non-uniform longitudinal sampling design, with data collected from each infant at up to 3 randomized time points between birth and 6 months (Fig. S1). Detailed demographic information is provided in Table S1. For all analyses, infant age at each scan was corrected for gestational age at birth, calculated as the postnatal age minus 7*(40 - gestational age in weeks). This corrected age was used to define the six-month postnatal period, corresponding to an age range of 0 to 180 days.

Preprocessing

We performed standard fMRI preprocessing using the FMRIB Software Library (FSL v6.0, https://fsl.fmrib.ox.ac.uk/fsl/fslwiki/) and the Statistical Parametric Mapping (SPM12, http://www.fil.ion.ucl.ac.uk/spm/) toolboxes under the MATLAB 2020b environment. Firstly, the initial ten dummy scans with substantial signal changes were discarded. Following that, we corrected the distortion in the images using SBRef data with phase encoding blips in reverse. After distortion correction, we performed slice timing to correct for these slice-dependent delays. Head motion correction was performed followed by the slice timing to realign all the scans to the reference scan. To normalize the infant data to the standard adult Montreal Neurological Institute (MNI) space, we applied a two-step normalization procedure. We first warped the UNC-BCP 4D Infant Brain Template (https://www.nitrc.org/projects/uncbcp_4d_atlas/)(44) into the adult MNI space using the EPI template as the reference. Here, to mitigate introducing an age-specific bias into normalization, we chose the template with the age of 4 months which is the median of the age range of our infant dataset. After having the UNC-BCP template in the adult MNI space, we normalized the infant data using it as the reference. Finally, the normalized fMRI data were spatially smoothed using a Gaussian kernel featuring a 6 mm full width at half maximum (FWHM).

Estimating ICN from rsfMRI

Independent component analysis (ICA) is a widely adopted data-driven method for extracting ICNs from rsfMRI(45). As a data-driven multivariate approach, ICA decomposes fMRI signals into statistically independent components (ICs), revealing spatially coherent brain networks without requiring prior assumptions about time courses or regions of interest(46–48). Additionally, ICA effectively isolates meaningful brain activity from various sources of noise, including physiological artifacts such as cardiac and respiratory signals, improving the robustness of ICN estimation (45). ICA has been extensively applied in large cohorts and across diverse imaging modalities, demonstrating its capability to uncover brain functional networks and connectivity features where conventional regression-based methods often fall short(49–51). Recent advances in infant neuroimaging have utilized ICA to identify networks during the first year of life that closely resemble those seen in adults(9, 11, 15, 17, 29) highlighting its utility in studying the maturation of brain connectivity.

#### *burst Independent Component Analysis (burstICA)*

The proposed burst independent component analysis (burstICA) method introduces a novel blind ICA approach designed to generate ICNs at multiple spatial scales, effectively mitigating the need to select a single optimal model order. The analysis pipeline, as illustrated in

Fig. S2, applies burstICA to 126 fMRI scans using sequential model orders ranging from 2 to 225, with a step size of 1.

The pre-processed resting-state fMRI (rsfMRI) data underwent a two-stage Principal Component Analysis (PCA) to effectively reduce dimensionality and optimize memory usage. The first stage involves a subject-specific spatial PCA, which normalizes the data to ensure that each subject contributes comparably to the shared subspace. This is mathematically represented as

$$X_{m\,(L \times V)} = F_{m\,(L \times K)}^{-1}\, Y_{m\,(K \times V)} \quad \text{(S1)}$$

where $X_{(C \times V)}$ is the reduced-dimensionality data for subject $m$, $F^{-1}$ is the reduction matrix obtained from PCA, $L$ is the reduced dimensionality of time points, $K$ is the original number of time points, $V$ represents the number of voxels, and $Y_m$ is the preprocessed data matrix for subject $m$. This PCA step reduces the dimensionality of the input time points from $K$ to $L$, effectively capturing the most significant spatial patterns while reducing the computational load for subsequent analyses. After performing the subject-specific PCA, the principal components derived for each subject were concatenated along the time dimension, creating a combined dataset that retains the key features of each subject's brain activity, facilitating group-level analysis. Subsequently, a group-level spatial PCA was applied to the concatenated matrix of subject-level principal components to further reduce the dimensionality and identify patterns common across all subjects. This process is described by

$$X_{(C \times V)} = G^{-1}{}_{(C \times LM)} \begin{bmatrix} F_1^{-1} Y_i \\ \vdots \\ F_M^{-1} Y_i \end{bmatrix}_{(LM \times V)} \quad \text{(S2)}$$

Here $G^{-1}$ is the group-level reduction matrix, $C$ is the number of components determined by the chosen model order, $M$ is the total number of subjects, and $LM$ is the product of the reduced dimensionality $L$ and the number of subjects $M$. This group-level PCA step provides a common subspace that reflects the shared structure across all subjects, preparing the data for the next step of ICA. The group-level principal components derived from the previous step were then used as input for spatial ICA to evaluate group independent components. This is mathematically expressed as

$$X_{(C \times V)} = A_{(C \times C)} S_{(C \times V)} \quad \text{(S3)}$$

In this equation, $A$ represents the mixing matrix that combines the independent components, and $S$ represents the estimated independent sources. The objective of ICA is to estimate an unmixing matrix, $W$ that can transform the mixed signals back into independent sources, described by

$$Y_{(C \times V)} = W_{(C \times C)} X_{(C \times V)} \quad \text{(S4)}$$

where $Y$ is the output representing a good approximation of the independent latent sources $S$. The Infomax principle, central to ICA, guides the optimization of the unmixing matrix $W$. According to this principle, the mutual information between the input signals $X$ and the output signals $Y$ should be maximized, which is mathematically represented as

$$I(X, Y) = H(Y) - H(Y|X) \quad \text{(S5)}$$

Here, $H(Y)$ is the entropy of the output signals, representing the amount of uncertainty or information content, and $H(Y|X)$ is the conditional entropy, minimized in this context. Maximizing the mutual information $I(X, Y)$ is equivalent to maximizing the entropy $H(Y)$ of the output signals, encouraging the components to be as statistically independent as possible, described by

$$H(Y) = H(y_1, y_2, \dots, y_c) \quad \text{(S6)}$$

Assuming that the components are independent, which aligns with the ICA goal, the entropy of the output signals can be rewritten as:

$$H(Y) = \sum_{c=1}^{C} H(y_c) \quad \text{(S7)}$$

The Infomax ICA algorithm iteratively adjusts $W$ to maximize the entropy of the outputs $Y$, thereby enhancing the separation of mixed signals into independent components. By optimizing $W$ using the Infomax criterion, the ICA algorithm effectively separates the mixed signals into their group-level ICs. To refine this process, the algorithm iterates through group-level PCA and Infomax ICA for each model order $C$, where $C$ varies sequentially, $(C = a, a + 1, , \dots, , k - 1, k)$. For the purpose of our analysis, we used the values of $C$ from 2 to 225 covering a broad range of model orders, from minimalistic to the highest order used in recent studies (4). This iterative process allows for the exploration of various levels of data granularity, ensuring a thorough analysis across multiple scales. The aggregated group-level ICs were matched with the multi-scale NeuroMark 2.2 template to identify the most similar among all the ICs using Pearson's correlation.

The component matching led to the identification of the top 105 best-matched ICNs from infant dataset, providing a robust set of ICNs that align closely with the NeuroMark 2.2 template.

Finally, to obtain subject-level ICN, the back reconstruction step was conducted using spatially constrained ICA that employs a multi-objective function optimization approach - Multi-Objective Optimization Independent Component Analysis with Reference (MOO-ICAR), which is detailed in NeuroMark framework(52) section below. This method reconstructs subject-specific networks based on the 105 selected group-level ICNs, generating both spatial maps and time courses for each of the 126 subjects. The resulting time courses were then utilized to evaluate functional network connectivity, offering insights into the functional organization of the infant brain.

#### *NeuroMark framework: A Reference-Informed Approach*

In this section, we introduce a reference-informed approach using the NeuroMark framework. Specifically, we employed the Multi-Objective Optimization Independent Component Analysis with Reference (MOO-ICAR), a spatially constrained ICA method for estimating ICNs, utilizing the NeuroMark 2.2 template as a reference. This approach allows us to evaluate both the accuracy of ICN estimation in infants and the practical feasibility of NeuroMark framework. We implemented MOO-ICAR using the GIFT toolbox(43) using the multi-scale NeuroMark 2.2 template (available at https://trendscenter.org/data/) as the spatial prior, facilitating the estimation of corresponding subject-specific ICNs. The details of this methodology are illustrated in Fig. S3.

MOO-ICAR is a spatially constrained ICA that employs a predefined template or reference to facilitate the identification of components, such that it aligns with the spatial characteristics of the reference. This approach aims to identify corresponding subject-specific ICNs across different datasets or populations based on a reference, usually obtained from a large dataset or a priori knowledge (4, 28). Mathematically, MOO-ICAR maximizes two objective functions to estimate each subject-specific network corresponding to a given template(28). The first objective functions, $J(S_I^k)$ aims to optimize the independence of the networks and the second objective function, $F(S_I^k)$ aims to optimize the correspondence between each subject-specific network and the template. Equation 1 summarizes how the $I^{th}$ network can be estimated for the $k^{th}$ subject using the network template $S_I$ as guidance.

$$max \begin{cases} J(S_I^k) = \{E[G(S_I^k)] - E[G(v)]\}^2 \\ F(S_I^k) = E[S_I S_I^k] \end{cases} \quad s.t. \left\|w_I^k\right\| = 1 \qquad \text{(S8)}$$

In this formulation, $S_I$ represents the $I^{th}$ network of the template and $S_I^k = (w_I^k)^T . X^k$ signifies the estimated $I^{th}$ network of the $k^{th}$ subject. $X^k$ is the whitened fMRI data matrix of the $k^{th}$ subject and $w_I^k$ represents the unmixing column vector that needs to be solved within the optimization functions. The first function $J(S_I^k)$ uses negentropy to optimize the independence measure of $S_I^k$. Here, $v$ denotes a Gaussian variable with mean zero and unit variance. The function $G(\bullet)$ is a nonquadratic function, and $E[\bullet]$ denotes the expectation of the variable. The second function $F(S_I^k)$ measured the correspondence between the template network ($S_I$) and subject network ($S_I^k$). The two objective functions are merged using a linear weighted sum with weights set at 0.5 to solve the optimization problem. Therefore, MOO-ICAR yields $N$ subject-specific networks corresponding to $N$ network templates, along with the relevant time courses for each scan.

Using this approach, we estimated 105 subject-specific multi-scale ICNs for each scan, including their spatial maps and time courses. To assess the correspondence between the estimated ICNs and the NeuroMark 2.2 template, we computed Pearson's correlation coefficients using the average spatial maps derived from the estimated ICNs. We then evaluated the unique information contained within each ICN estimated by the NeuroMark framework for each subject. For each target ICN from the NeuroMark 2.2 template, we identified a comparison ICN that had the second highest spatial correlation with the target ICN, with the estimated ICN itself being the highest and the network of interest. Using these ICNs, we then extracted their equivalents from the infant dataset. We computed Pearson's correlation with the target ICN for the network of interest and the comparison ICNs from the dataset. This analysis allowed us to evaluate the preservation of network-specific information across subjects. The time courses were further used to generate Functional Network Connectivity (FNC) by correlating time courses obtained for each ICN.

Functional Network Connectivity

Prior to FNC computation, ICN time courses were cleaned to reduce motion-related and physiological artifacts. Cleaning involved z-scoring, regression of 6 head motion parameters and their temporal derivatives, removal of low-frequency polynomial trends (up to cubic), despiking, and zero-phase band-pass filtering (0.01–0.15 Hz).

Static FNC was then computed as pairwise Pearson correlations between the cleaned time courses of all 105 scan-specific ICNs, producing a symmetric 105 × 105 connectivity matrix for each scan. Group-average matrices were compared to an adult reference FNC from the UK Biobank(23) to assess developmental correspondence.

**Fig. S1.**

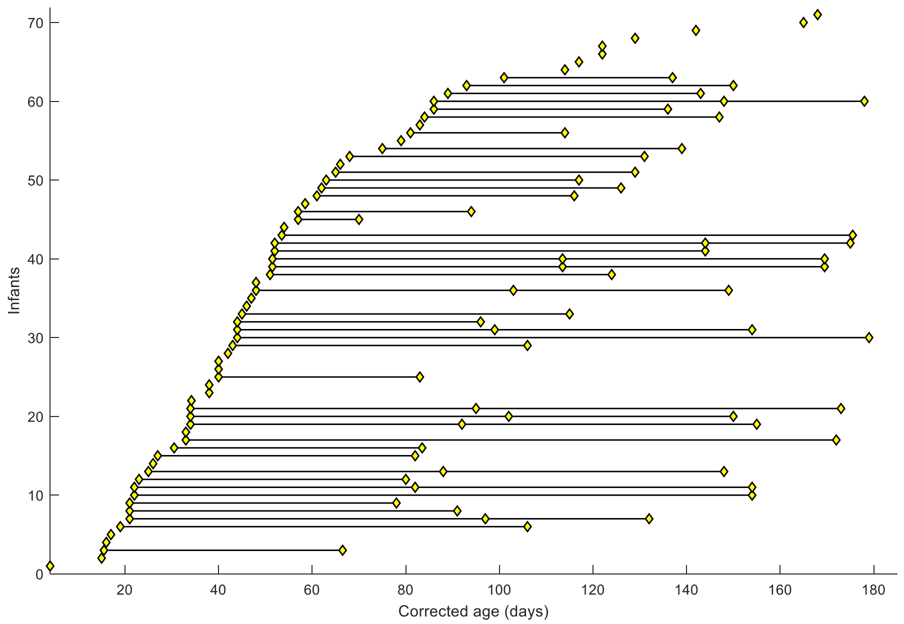


Fig. S1. **Included scans for all infant participants.** Yellow diamonds represent a scan, with lines connecting scans collected from the same participant

**Fig. S2.**

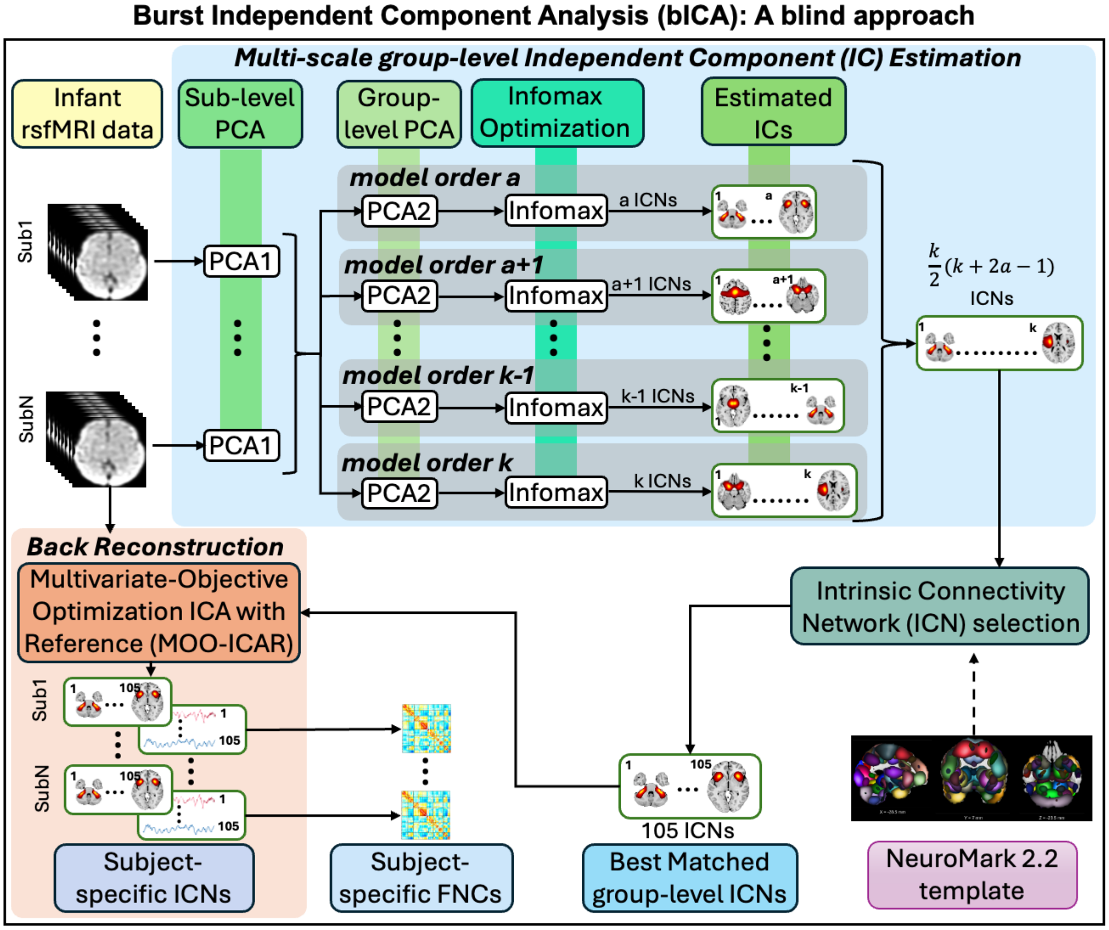


Fig. S2. **A pipeline for burst independent component analysis (burstICA)**, a blind approach involving estimation of independent components (ICs), ICN selection, and back-reconstruction. The method initiates with subject-level PCA, followed by group-level PCA and Infomax optimization, iterated across multiple model orders to generate a comprehensive set of multi-scale ICs, from which the top 105 ICs were selected based on Pearson's correlation with the NeuroMark 2.2 template. These ICNs served as references for back-reconstruction, specifically MOO-ICAR, to generate subject-level spatial maps and time courses. Subsequently, the time courses were used to generate functional network connectivity (FNC).

**Fig. S3.**

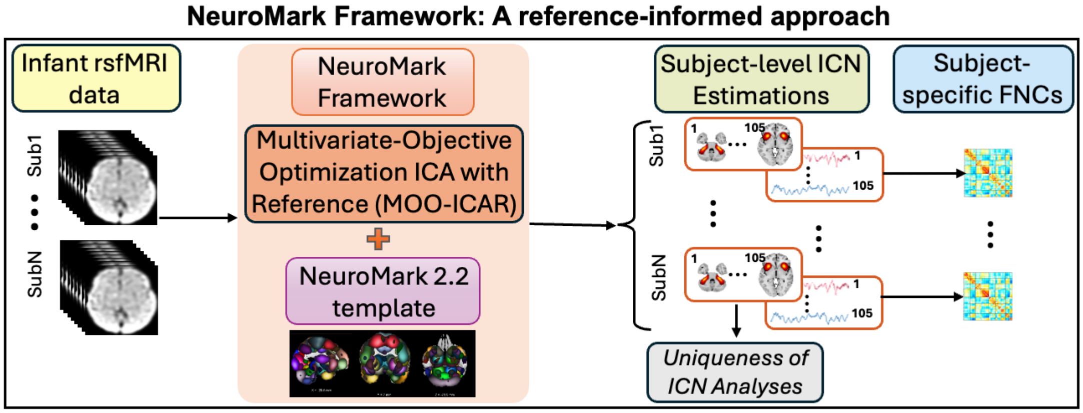


Fig. S3. **A pipeline for NeuroMark framework**, a reference-informed approach that uses MOO-ICAR method along with NeuroMark 2.2 Template as a reference to generate precisely 105 subject-level ICN estimations from the infant dataset. The spatial maps are further used to analyze the unique information in each ICN, and the time courses are used to evaluate functional network connectivity (FNC).

**Table S1.**

Table S1. **Demographics of participant sample.** For information categories with a participant number less than the total sample, the N is specified next to the category title.

| | (N = 71) |
|---|---|
| Infant Sex | 30f, 41m |
| Gestational Age at Birth, mean (SD) | 39.1wks (1.27) |
| Race (N=64) | |
| Black | 7.8% |
| White | 87.5% |
| More than one race | 4.7% |
| Maternal Education (N=60) | |
| High School | 1.7% |
| Trade/Vocational Training | 1.7% |
| College Courses | 5.0% |
| Associate's Degree | 3.3% |
| College Degree | 36.7% |
| Graduate Degree | 51.6% |
| Household Income (N=61) | |
| < $40000 | 4.9% |
| $40000-$80000 | 19.7% |
| $80001-$100000 | 23.0% |
| $100001-$150000 | 26.2% |
| > $150001 | 26.2% |
